# Beyond Linear Neural Envelope Tracking: A Mutual Information Approach

**DOI:** 10.1101/2022.08.11.503600

**Authors:** Pieter De Clercq, Jonas Vanthornhout, Maaike Vandermosten, Tom Francart

## Abstract

The human brain tracks the temporal envelope of speech, which contains essential cues for speech understanding. Linear models are the most common tool to study neural envelope tracking. However, information on how speech is processed can be lost since nonlinear relations are precluded. As an alternative, mutual information (MI) analysis can detect both linear and nonlinear relations. Yet, several different approaches to calculating MI are applied without consensus on which approach to use. Furthermore, the added value of nonlinear techniques remains a subject of debate in the field. To resolve this, we applied linear and MI analyses to electroencephalography (EEG) data of participants listening to continuous speech. Comparing the different MI approaches, we conclude that results are most reliable and robust using the Gaussian copula approach, which first transforms the data to standard Gaussians. With this approach, the MI analysis is a valid technique for studying neural envelope tracking. Like linear models, it allows spatial and temporal interpretations of speech processing, peak latency analyses, and applications to multiple EEG channels combined. Finally, we demonstrate that the MI analysis can detect nonlinear components on the single-subject level, beyond the limits of linear models. We conclude that the MI analysis is a more informative tool for studying neural envelope tracking.

**Significance statement:** In the present study, we addressed key methodological considerations for MI applications. Traditional MI methodologies require the estimation of a probability distribution at first. We show that this step can introduce a bias in the results and, consequently, severely impact interpretations. As an alternative, we propose using the parametric Gaussian copula method, which we demonstrated to be robust against biases. Second, using the parametric MI analysis, we show that there is nonlinear variance in the EEG data that the envelope of speech can explain at the single-subject level, proving its added value to neural envelope tracking. We conclude that the MI analysis is a statistically more powerful tool for studying neural envelope tracking than linear models. In addition, it retains spatial and temporal characteristics of speech processing which are lost when using more complex deep neural networks.

## 1 Introduction

When listening to natural speech, the human brain tracks certain stimulus features required for speech understanding. The speech envelope is the feature investigated most intensively and explains most of the variance observed in the neural response. The envelope contains the slow-varying temporal modulations in the speech signal. The low frequencies (< 8 Hz) are often the main interest for researchers, as they contain the syllabic, word, and phrase rates, which are crucial for speech understanding (Peelle and Davis, 2012). There is abundant evidence from studies supporting this claim by showing decreased envelope tracking as the level of intelligibility of the speech signal decreases (Ding and Simon, 2013; Etard and Reichenbach, 2019; Vanthornhout et al., 2021). This is in line with behavioral results showing that listeners can understand speech based on the low-frequency envelope only (Shannon et al., 1995).

The neural response to the speech envelope can be modeled in several ways. The most popular approach is to use linear models. In the linear backward model, the speech envelope is reconstructed from a weighted sum of the EEG channels and their time-shifted version to account for neural processing delays. The reconstruction accuracy is obtained by correlating the reconstructed and the original envelope and is considered a measure of how well speech is tracked by the brain (i.e., envelope tracking). The backward model is a powerful tool to decode the envelope by optimally combining the individual channels. On the other hand, the model weights are rather hard to interpret (Haufe et al., 2014). As an alternative, the linear forward model is an excellent tool for investigating the spatial and temporal properties of the neural response. In the forward model, the neural signal of each channel individually is predicted from a weighted sum of the speech envelope at different time delays. The resulting weights form the temporal response function (TRF), with the magnitude of the weights reflecting the response amplitude of the neural signal at different response latencies. The TRF thus reflects how the brain processes speech over time. A measure of envelope tracking in the forward model can be obtained by correlating the predicted with the original neural response.

The linear forward model has a natural advantage over more complex models (such as neural networks, Accou et al. (2021)) since the TRF’s are interpretable temporally and spatially. Yet, information on how speech is processed can be lost, as the brain (including the auditory cortex) is known to employ nonlinear relationships (Ahrens et al., 2008; Sahani and Linden, 2003), which linear models cannot capture. Deep neural networks can model nonlinear relationships, but they are black boxes and preclude spatio-temporal interpretations. A compromise between interpretability and deriving nonlinear relationships could however be made by relying on an information-theoretic approach. Mutual information (MI) analyses are a powerful tool to investigate statistical dependency between two variables. Both linear and nonlinear relationships influence MI, with higher MI values reflecting higher dependencies between variables. A temporal pattern can be obtained by shifting one variable in time and calculating the MI at each timestep, forming the ‘Temporal Mutual Information Function’ (TMIF). Zan et al. (2020) recently applied this procedure to a group study comparing envelope tracking in healthy younger and older adults. The TMIF showed response components at specific latencies and spatial patterns similar to the TRF. In addition, the TMIF had higher statistical power compared to the TRF, as it found statistically robust group differences that the TRF did not observe in a previous analysis of the same data (Brodbeck et al., 2018).

In sum, the advantage of MI is that it can find statistical effects beyond linear relationships while retaining spatial and temporal interpretations. For these reasons, MI analyses are gradually becoming a more popular approach to studying neural envelope tracking (Chalas et al., 2022; Coopmans et al., 2022; Daube et al., 2019; Giordano et al., 2017; Kaufeld et al., 2020; Keitel et al., 2018; Keshavarzi et al., 2021; Perez et al., 2022; Pfeffer et al., 2022). However, the field currently lacks a proper comparison between the different methodologies that exist to derive MI. Moreover, results obtained from an MI analysis have not yet been validated against more extensively documented linear models. Finally, it remains unclear to what extent this nonlinear approach provides an added value over linear models. Power and Reilly (2011) reported marginal improvements for a nonlinear model (quadratic TRF’s), but they used acoustic white noise as the stimulus. For speech stimuli, electrocorticography studies demonstrated that nonlinear models (deep neural nets) outperformed linear models (Akbari et al., 2019; Yang et al., 2015). Yet, few neural envelope tracking papers have investigated the added value of nonlinear techniques over linear models using noninvasive recordings (EEG, magnetoencephalography (MEG)). It further remains unclear whether the MI analysis is sensitive to capturing these relations. The present paper aims to resolve these open questions.

We first introduce the different methodologies that exist to calculate MI. Next, we show that certain MI calculations and parameter values can significantly impact interpreting experimental results. We then compare the MI analysis to the linear forward and backward model. Finally, we discuss the extent to which the MI analysis, as a nonlinear approach to neural envelope tracking, can provide an added value over linear models.

### 1.1 Calculating Mutual Information

MI has its grounds in Shannon’s Information Theory (Shannon, 1948). Formally, MI quantifies the amount of uncertainty (i.e., variability) of one random variable reduced by observing another random variable. The amount of uncertainty is referred to as entropy, often expressed in bits (binary logarithm):

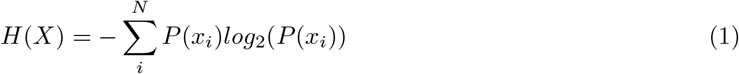

with *H*(*X*) equal to the entropy of the marginal distribution of variable *X* (e.g., the speech envelope), and *P*(*x_i_*) the probability of observing value *x_i_* with N possible values. The entropy of a second variable, *Y* (e.g., the neural response of a single channel), is computed according to the same formula. The MI of two variables can then be expressed as follows:

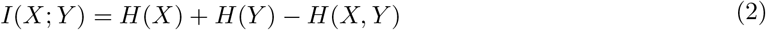

with

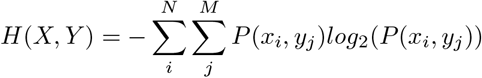

The MI between *X* and *Y*, *I*(*X*; *Y*), is thus equal to the sum of the entropy of the marginal distributions subtracted by the entropy of the joint distribution (i.e., the joint entropy *H*(*X*,*Y*)). MI quantifies the statistical dependency between *X* and *Y*, and, unlike a Pearson correlation, it measures both linear and nonlinear relationships. Temporal information of MI (i.e., the TMIF) in the context of envelope tracking is obtained by sliding the neural response in a channel (*Y*) in function of the envelope (*X*) for a certain integration window and calculating the MI at each timestep.

The marginal and joint entropy definitions involve the explicit knowledge of the probability distributions. In practice, however, these are not known and have to be estimated from the data, which is a nontrivial task. We dissociate three different methodologies to calculate the probability values and discuss them below. Appendix A includes a more in-depth explanation of these different methodologies.

#### The histogram-based method

The most common approach to estimating probability values is based on the histogram-based algorithm. Here, the data are first partitioned into a set of discrete bins. The probability of observing a discrete bin value is then obtained by dividing the number of occurrences in a single bin by the total number of data points. The probability values can then be plugged into the marginal and joint entropy formulas.

Binning most commonly occurs according to the equipartition principle, where bins have equal occupancy (e.g., if using 10 bins, each bin contains 10% of the data). An important consideration is how many bins one should choose for constructing the histogram. In theory, one should not have too few bins, otherwise, there will be little sensitivity to the distribution, and the relationship between two variables can be lost. On the other hand, if one has too many bins, estimation errors for the joint distribution can be made hence generality is lost. The latter case can be seen as a form of overfitting, as the distribution will become sensitive to noise components in the data. Therefore, the bin parameter can induce either an under-(too few bins) or overestimation (too many bins) to the true MI. Previous neural tracking studies have chosen the bin parameter rather arbitrarily with little justification or empirical data to support their choice (Kaufeld et al., 2020; Zan et al., 2020).

#### Continuous alternatives

Instead of partitioning the data into discrete bins, several continuous alternatives exist to estimate the probability distribution. In the Kernel Density Estimation (KDE) approach, each datapoint is convolved with a (most commonly Gaussian) kernel. The probability density function is then obtained by summing all kernel values and normalizing so the integral equals one. In the KDE approach, researchers need to pre-specify the width of the kernel (i.e., the bandwidth), which is the equivalent critical choice to the number of bins in the histogram-based approach (Venelli, 2010). The bandwidth parameter determines the smoothness of the obtained probability distribution. A bandwidth that is too large leads to over-smoothing, hence specificity is lost, while a bandwidth that is too small leads to under-smoothing, hence generality is lost. A recent neural envelope tracking paper calculated MI based on KDE estimations (Keshavarzi et al., 2021), but the bandwidth parameter was not specified.

Correctly choosing the number of bins or the bandwidth parameter is critical for constructing the probability distribution. Several rules of thumb exist, such as the Freedman-Diaconis rule (Freedlan and Diaconis, 1981) for histogram-based distributions and the Silverman’s rule (Silverman, 1986) for KDE estimations. Cross-validation approaches have also been proposed in the past (Sain et al., 1994). With these approaches, the obtained parameter value is data-specific. Yet, it is crucial to fix this parameter value to enable comparisons across participants and channels since different parameter values introduce different baseline MI values (Vinh et al., 2009). An alternative approach that bypasses the need to set the bins or bandwidth parameter is to assume that the data are normally distributed. Under this assumption, the MI between two Gaussian variables is determined based on the covariance matrices:

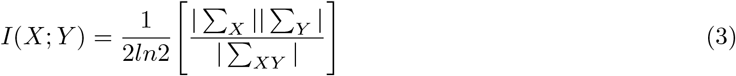

where ∑_*X*_ and ∑_*Y*_ are the covariance matrices of variables *X* and *Y*, and ∑_*XY*_ is the covariance matrix for the joint variable. Straight lines indicate that the determinant of the covariance matrices is calculated. The resulting algorithm does not require any parameter values. However, we must rely on the strong assumption that the data are normally distributed.

#### Gaussian copula method

Recently, Ince et al. (2017) introduced the concept of Gaussian copula MI for neuroimaging research. The method uses the Gaussian definition of MI but transforms the data so it meets the normality assumption. The procedure first implies that values for the individual variables are ranked on a scale from 0 to 1, obtaining the cumulative density functions (CDF’s). By computing the inverse standard normal CDF of the variables separately, the data distributions are transformed to perfect standard Gaussians. With these steps, the statistical relationship between both variables (i.e., the ‘copula’) is preserved. After data transformations to perfect standard Gaussians, the parametric MI estimate in Eq. (3) can be applied. Crucial, the MI in this method provides a lower bound to the true MI, i.e., erroneous high values cannot occur. This is because the joint Gaussian distribution has the maximum entropy for a variable with a given mean and covariance matrix (Cover, 1999). For a more in-depth explanation, we refer to Appendix A. Over recent years, the Gaussian copula MI has been applied in numerous neural envelope tracking papers (Coopmans et al., 2022; Daube et al., 2019; Giordano et al., 2017; Perez et al., 2022).

The Gaussian copula has a particular advantage over the other MI derivations in multidimensional spaces. While the histogram- and KDE-based approaches severely suffer from the curse of dimensionality and computational complexity, the Gaussian copula method can efficiently handle multidimensional variables. After transforming each individual dimension to a standard Gaussian, the formula in Eq. (3) can be applied. For the context of neural envelope tracking, this may open perspectives to calculating the MI between the speech envelope and multiple channels combined:

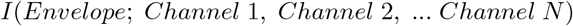

This could be considered an information-theoretic analog to the linear backward model.

#### Present study

In the present study, we compare the Gaussian copula approach with the probability density estimation approach in the context of neural envelope tracking. We restrict the comparison to the most popular approach, i.e., the histogram-based method, to demonstrate the effect of arbitrarily setting a parameter (i.e., the number of bins) to estimate the probability distribution. Next, we validate the most suitable MI technique against the linear forward and backward model. To this end, we apply the measures to 2 datasets containing EEG data of participants listening to continuous speech. The first dataset includes a large group study of healthy young participants listening to a natural story. With this dataset, we compare the temporal and spatial patterns of the TMIF and TRF in-depth, discuss the properties of multivariate MI in relation to the linear backward model, and investigate whether non-linear components are present in the neural response to the speech envelope. The second dataset includes normal hearing younger and older adults listening to continuous speech to demonstrate the application of MI in a group study and highlight certain interesting differences between MI and its linear counterpart.

## 2 Methods

We discuss two datasets where we applied MI measures and linear models to EEG data. All EEG data were recorded in a soundproof, electromagnetically shielded booth using a 64-channel BioSemi ActiveTwo system (Amsterdam, Netherlands) at a sampling frequency of 8,192 Hz. Stimuli were presented through ER-3A insert earphones (Etymotic Research Inc, IL, USA) using the software platform APEX (Francart et al., 2008).

### 2.1 Dataset 1: Natural Story Listening

The first dataset was used to characterize the properties of MI and discuss them in relation to linear models. The data, described in previous studies (Accou et al., 2021), comprise a large group of 64 healthy young participants (49 female, mean age = 20.4 years, SD = 1.87 years) listening to the first part of the story ‘De Kleine Zeemeermin’ (15 minutes, written by H. C. Andersen and narrated by Katrien Devos, female) in silence. All participants signed informed consent and had normal hearing, verified using pure tone audiometry (threshold below 20 dB hearing level for octave frequencies between 125 and 8,000 Hz). Stimuli were presented bilaterally at 62 dBA.

### 2.2 Dataset 2: Healthy Aging

The second dataset consists of 54 healthy normal hearing participants (41 female) of varying ages listening to the story ‘Milan’ (written and narrated by Stijn Vranken) while their EEG was recorded. The participants were divided into 3 groups: young (17-37 yo, N = 11), middle-aged (42-60 yo, N = 32) and old (61-82 yo, N = 11). We refer to the original paper of Decruy et al. (2019) for a detailed description of the demographics and the methods. The data were recently reanalyzed with the forward modeling approach showing higher neural tracking for younger adults compared to elderly (Gillis et al., 2022). Here, we apply MI to demonstrate certain interesting properties of this method for group comparisons.

All participants signed informed consent and had normal hearing, verified using pure tone audiometry (thresholds lower or equal to 30 dB hearing level for octave frequencies between 125 and 4,000 Hz). The stimuli were presented unilaterally (to the right ear, except for one participant) at 55 dBA.

### 2.3 Signal Processing

#### Envelope extraction

For both datasets, the speech envelope was extracted using a gammatone filter bank (Søndergaard et al., 2012), with 28 channels spaced by 1 equivalent rectangular bandwidth and center frequencies from 50 Hz until 5000 Hz. The envelopes were extracted from each subband by taking each sample’s absolute value and raising it to the power of 0.6. The resulting 28 subband envelopes were averaged to obtain one single envelope. The envelope was then downsampled to 512 Hz to decrease processing time. Next, the envelope was highpass-filtered at 0.5 Hz (transition band 0.45-0.5 Hz) and lowpass-filtered at 8 Hz (transition band 8-8.8 Hz) using a Least Squares filter of order 2000 and compensated for the group delay. After filtering, the envelope was normalized and further downsampled to 128 Hz.

#### EEG processing

The EEG signals were first downsampled to 512 Hz to decrease processing time. Eye-blink artifacts were removed using a multi-channel Wiener filter (Somers et al., 2018) and the EEG was referenced to common average. The data were then highpass-filtered at 0.5 Hz and lowpass-filtered at 8 Hz using the same Least Squares filter used in the envelope extraction method and compensated for the group delay. Next, normalization and additional downsampling to 128 Hz were applied.

#### Linear forward model

In the linear forward model, the EEG is predicted as a function of the speech envelope. This approach estimates the Temporal Response Function (TRF), a kernel that describes the neural response at different latencies to the speech envelope. We used the Eelbrain toolbox (Brodbeck, 2020) to estimate the TRF for each EEG channel separately using the boosting algorithm (David et al., 2007). To avoid overfitting, a 10-fold cross-validation was used (10 equally long folds, 8 for training, 1 for validation and 1 for testing). The TRF covered an integration window from −200 to 600 ms, with a 50 ms Hamming window basis. Training stopped when the mean squared error on the validation data stopped decreasing. Next, the TRFs were used to predict the EEG-response in the test fold by convolving it with the speech envelope. We then calculated the Spearman correlation between the predicted EEG and the actual EEG per channel, yielding a measure of neural tracking. The TRFs were averaged across all folds to interpret the temporal and spatial patterns.

#### Linear backward model

In the linear backward model, the speech envelope is reconstructed from the neural data. A linear decoder attributes weights to each EEG channel and a time-shifted response of each channel. The reconstruction of the speech envelope *ŝ*(*t*) is obtained through:

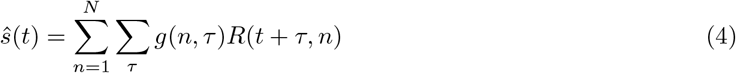

where *g* is the linear decoder as a function of the number of electrodes *N* and the integration window *τ*, (−200 to 600 ms). Matrix *R* contains the shifted neural responses of all channels (with time index *t*). The decoder was calculated by solving *g* = (*RR^t^*)^−1^(*Rs^t^*) with *s* the speech envelope. To prevent overfitting, ridge regularization and 10-fold cross-validation (10 equally long folds, 9 used to train the decoder and 1 used for testing) were applied. In the test fold, the reconstructed envelope was correlated with the actual envelope yielding a measure of neural tracking. Results were averaged across all folds. For a more in-depth explanation of linear models and ridge regression, we refer to Crosse et al. (2021).

#### Mutual information Analyses

The MI between the envelope and the EEG was calculated using the histogram-based and Gaussian copula methods. The MI in the histogram-based approach is obtained by applying Eq. (2) in Section 1.1. Histograms were constructed according to the equipartition principle, where bins have varying widths yet equal observation probabilities (hence, we obtain uniform distributions). We partitioned the data into several numbers of bins to demonstrate the effect of this parameter on the results. The Gaussian copula MI was calculated by applying the summarized data transformations in Section 1.1 and the Gaussian definition of MI (see Eq. 3). The analyses for the Gaussian copula MI were performed with the ‘GCMI’-toolbox (Ince et al., 2017). For both MI measures, the temporal evolution of the MI (i.e., the Temporal Mutual Information Function, TMIF) is obtained by shifting the EEG relative to the speech envelope from 200 ms prestimulus to 600 ms poststimulus and recalculating the MI at each sample.

The MI between all individual channels and the envelope was calculated to derive spatial interpretations. This measure will be compared to the linear forward model. The Gaussian copula approach further allows calculating the MI between the speech envelope and all channels combined by computing the covariance matrices of multidimensional data. This multivariate MI will be compared to the analogous linear counterpart, i.e., the backward model, where the envelope is reconstructed as a weighted sum of multiple channels.

#### Significance level

All methods were subject to permutation testing to quantify the meaningfulness of the derived values. We created stationary noise that matched the spectrum of the EEG responses. This was repeated 1000 times for each channel and each participant individually. For each repition, we calculated the MI, the prediction accuracy (linear forward model) or the reconstruction accuracy (linear backward model). The significance level was then determined as the 95th percentile of the neural tracking result of all repetitions.

For the multivariate MI, permutations were created from noise matching the spectrum of the envelope rather than noise matching the spectrum of the EEG. This was done to prevent dependencies between EEG channels from disappearing.

#### Test for nonlinear components

We investigated whether nonlinear relationships are captured by MI in the neural response to the speech envelope, both for individual channels and for the multivariate MI (all channels combined). To this end, we first applied linear models to remove all linear components in the data. The predicted EEG was subtracted from the actual EEG for the forward model. Next, the MI was calculated between the speech envelope and the residual EEG. For the multivariate MI, the linear backward model reconstructed the envelope, and this was subtracted from the actual envelope. The MI between the residual envelope and the 64-dimensional EEG data (64 channels) was then calculated. The results were compared to the significance level.

We used ordinary least squares without regularization to ensure that all linear components that the linear models could find were removed entirely from the data. Weights and correlations were then exactly equal to 0 (within machine precision) in a second linear fit, performed as a sanity check.

#### Channel selection

We will refer to a central and a parieto-occipital channel selection in the results section to plot the results in channels that exhibit the same polarity and that are known to contribute to auditory speech perception (Lesenfants et al., 2019). These respectively include channels [FC5, FC3, FC1, C5, C3, C1, FC6, FC4, FC2, C6, C4, C2, FCz, Cz] and [TP7, CP5, P5, P7, P9, PO7, PO3, O1, TP8, CP6, P6, P8, P10, PO8, PO4, O2, Iz, Oz,POz]. Since MI measures are strictly positive, we combine both clusters into a single large cluster. We will also refer to a control region (i.e., frontal channels) where no envelope tracking is expected. This control region includes channels [Fpz, Fp1, AF7, F7, Fp2, AF8, F8, AFz, AF3, AF4]. The channel selections are depicted in Supplementary Figure 1.

## 3 Results

### 3.1 Comparing Different Mutual Information Derivations

We first compare the properties of the histogram-based approach to the Gaussian copula approach for Dataset 1. We then compare MI to linear models in subsequent sections. Figure 1 provides an overview of the temporal and spatial patterns resulting from the different methods: Figure 1A displays the Temporal Mutual Information Function (TMIF) derived from the histogram-based approach with 10 bins, Figure 1B the Gaussian copula TMIF, and Figure 1C shows the Temporal Response Function (TRF) in the linear forward model.

**Figure 1.**
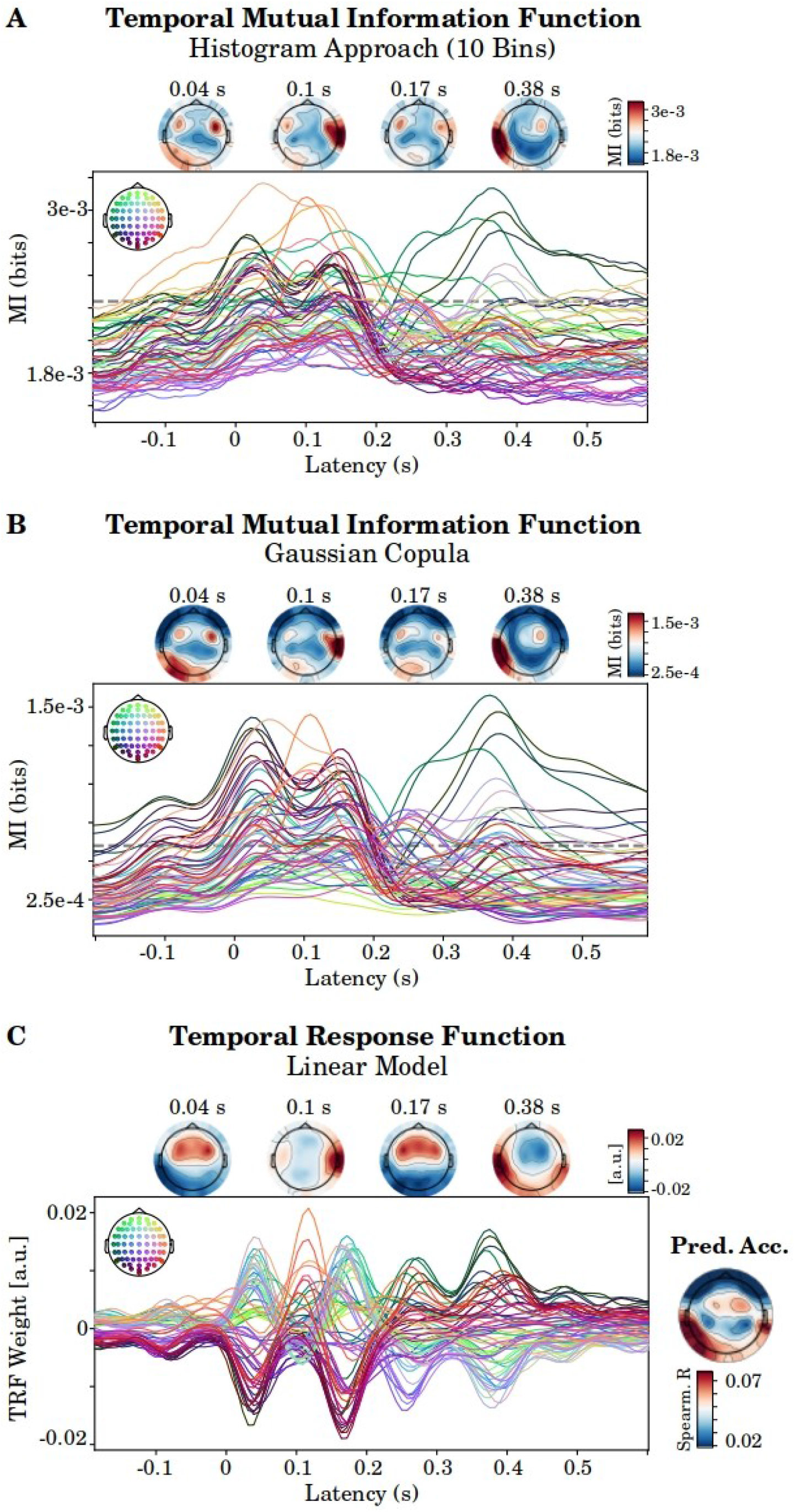
Overview of the different methodologies. **A.** The TMIF for the histogram-based approach with 10 bins. **B.** The TMIF for the Gaussian copula approach. **C.** The TRF in the linear forward model, and the topography of the prediction accuracy. Individual lines represent the group average per channel. The horizontal grey, dashed line in Panel A and B represents the 95th percentile of permutations.

#### The histogram-based approach introduces biases

##### 1. For different channels

The histogram-based approach requires a prespecified number of bins. Figure 2 illustrates the effect of this parameter value on the TMIF for Dataset 1. When taking 2 bins (Figure 2A), the variability across channels is minimal, but sensitivity to the full relationship is lost. For example, the left-lateralized temporal peak at 380 ms, present with 4/10 bins and present in the linear TRF, is largely suppressed. With 4 bins, specificity is preserved compared to the pattern that results from a larger number of bins (10, Figure 1A). But when the number of bins is too high (20), the spatial pattern changes drastically: the MI in frontal channels dramatically increases compared to the other channels (Figure 2C). Higher MI values for the frontal channels are even present before 0 ms, where the brain cannot yet process speech. We further observe that the topography of the significance level of frontal channels, obtained through calculating the MI between the envelope and spectrally matched EEG-shaped noise, is higher than the significance level of our channel selection (see Figure 2D), irrespective of the number of bins. The histogram-based method thus introduces a bias in the results: some channels systematically exhibit higher MI values than others by chance.

**Figure 2.**
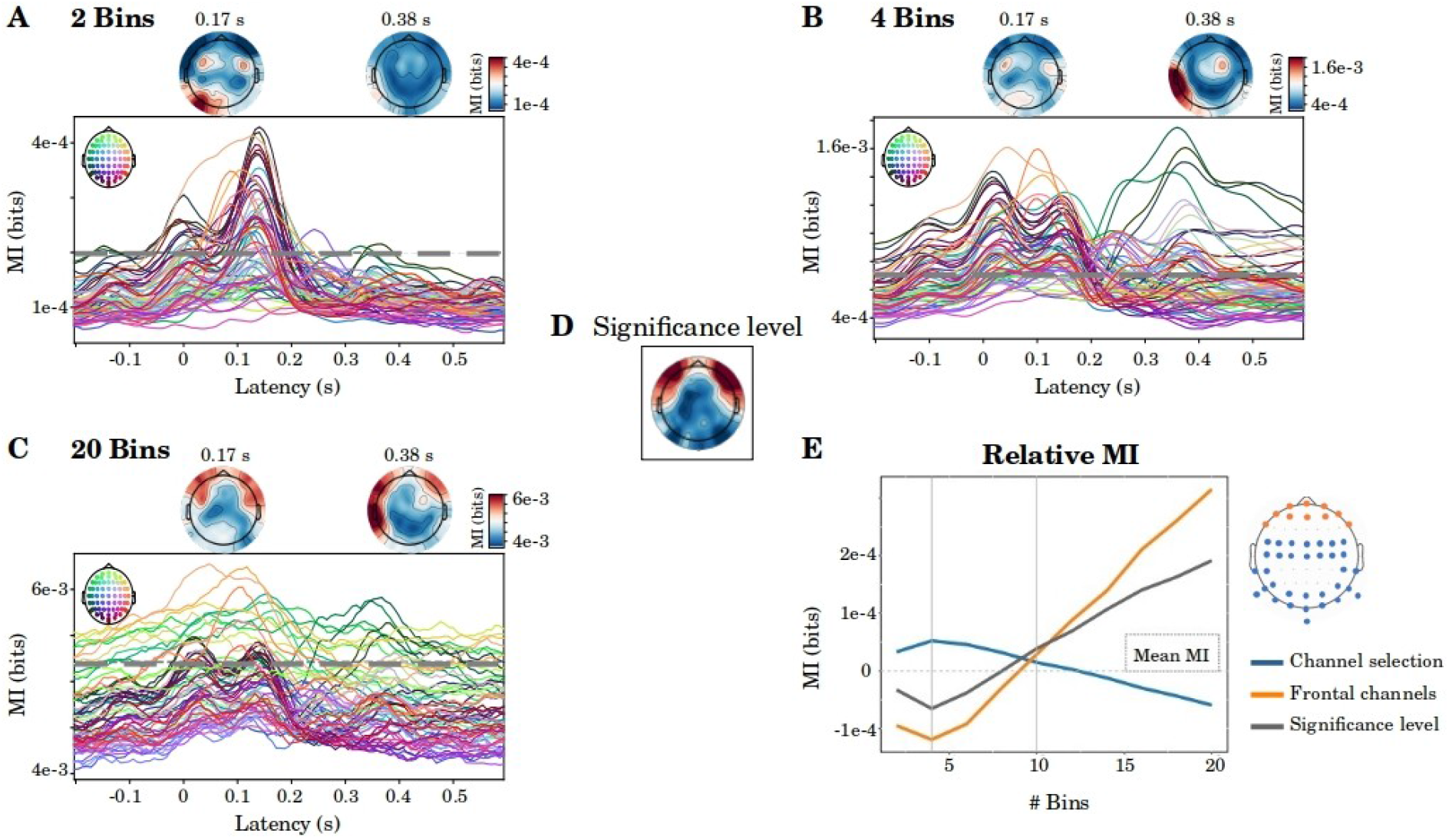
Impact of the number of bins in the histogram-based approach. Panel A, B and C depict the group average MI using 2, 4 and 20 bins, respectively. The dashed grey line indicates the significance level. **D.** Topography of the significance level. **E.** Relative MI values as a function function of the number of bins. This is obtained by calculating the average of our channel selection of interest (blue) and frontal channels (orange) in a 0 to 500 ms integration window, and subtracting the average with the global average of all channels. This demonstrates that the spatial distribution of MI changes as a function of the number of bins. The grey line represents the significance level, and shows that the envelope is no longer tracked significantly from 8 bins on (significance-level is higher than 0, which is the average across channels).

The issue of unequal baseline levels across channels has implications, for example, when interpreting the spatial distribution of MI. We demonstrate this in Figure 2E. The relative MI values for frontal channels and our channel selection were calculated by subtracting its average from the global average (all channels). We plot these relative measures as a function of the number of bins. For illustration, we interpret the case of 4 and 10 bins, two parameter values previously set in the field of envelope tracking (Gross et al., 2013; Zan et al., 2020). If 4 bins were selected, one might conclude that there is relatively higher neural tracking in our channel selection. Conversely, MI is higher for the frontal channels when using 10 bins. Thus, the number of bins that the researcher arbitrarily sets influences the spatial distribution of MI. Compared to significance-level, Figure 2E shows that from 8 bins on, the envelope is no longer tracked significantly for the mean of all channels (gray line crosses the mean MI value).

Unequal baseline MI across channels originates from differences in power in the lowest frequencies. We illustrate this bias with 30 seconds of EEG data of 1 participant in Supplementary Figure 2. Channels with relatively high power in the low frequencies (0.5-1 Hz) have a more slowly oscillating behavior and remain in the same bin for a more extended consecutive period of time (see Panels A and B). As a result, when no true relationship is present, the joint entropy with a random variable has a lower expected value: periods of high occurrences for certain bin combinations alternate with low to zero occurrences for other combinations in the joint distribution (Panel C). Lower joint entropy is directly related to higher MI (see Eq. (2)). Therefore, channels with high power in the low frequencies have a higher expected MI by chance. Across channels, we report a strong correlation between power in the low frequencies (0.5-1 Hz) and the significance level (Spearman’s r=0.75, p<0.001).

The MI based on the Gaussian copula, on the other hand, entails a transformation of the marginal distributions to standard Gaussians. The calculation of MI is then based on the covariance matrices. Hence, it does not require prespecified parameter values. For the Gaussian copula TMIF, we observe a pattern similar to the histogram-based approach, but with the additional benefit that the frontal channels, where no true relationship between the EEG and the envelope is expected, are effectively suppressed (see Figure 1B).

##### 2. For different stimuli

The histogram-based MI thus suffers from a bias induced by power in the low frequencies. Yet, this bias arises in the stimulus as well. We illustrate this with the 2 stimuli we consider in this paper: the story ‘De Kleine Zeemeermin’ (‘DKZ’) for Dataset 1, and ‘Milan’ for Dataset 2. The envelope of the latter story has higher power in the low frequencies, visualized by the power spectrum (Figure 3A). As a result, the latter story has higher baseline MI values (obtained through calculating the MI with EEG-shaped noise). This bias again exacerbates with a higher number of bins. The Gaussian copula MI, on the other hand, is much more robust to this issue: baseline neural tracking measures are approximately equal for both stimuli (Figure 3C).

**Figure 3.**
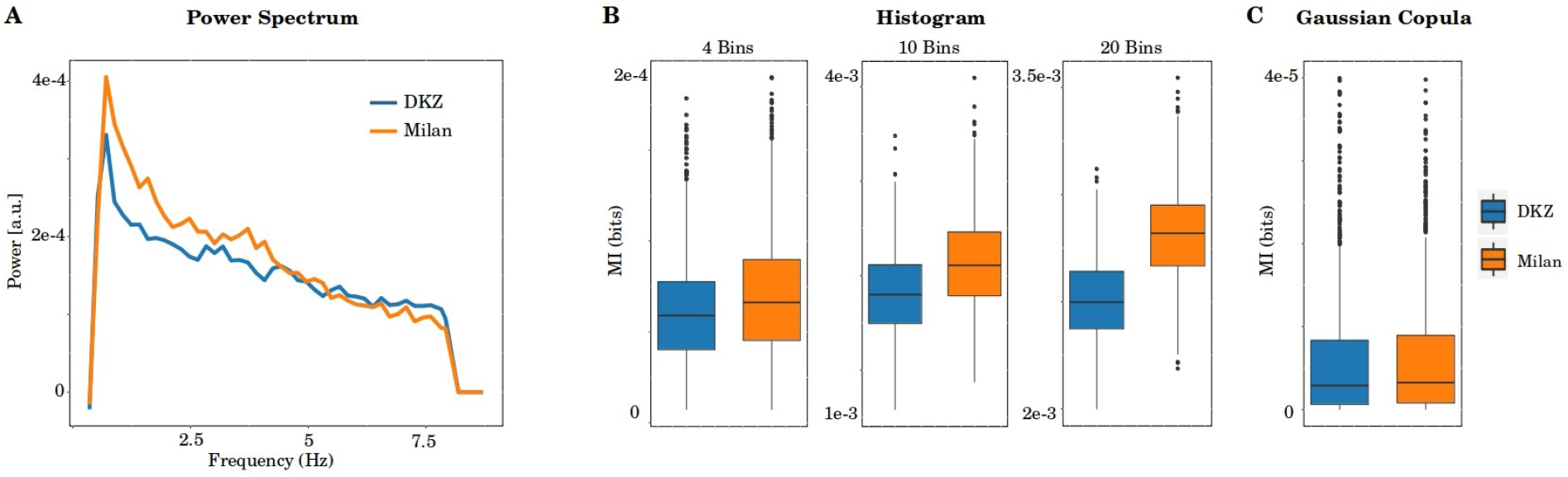
Bias in the results induced by the power spectrum of the stimulus. **A.** Power spectrum of the envelope for the story ‘De Kleine Zeemeermin’ (‘DKZ’) and ‘Milan’. **B.** Baseline MI values (MI between the envelope and 1000 resamplings of EEG-shaped noise) for the histogram-based approach with 4, 10 and 20 bins. **C.** Baseline MI values for the Gaussian copula approach.

### 3.2 Mutual Information in Relation to the Linear Forward Model

For the reasons listed above, namely that unequal baseline values naturally occur for different channels and stimuli, we no longer consider the MI based on the histogram. Instead, we focus on the robust measure of Gaussian copula MI. We now compare its properties to the well-documented linear forward model. First, we compare the temporal and spatial patterns of both measures (depicted in Figure 1). Second, we investigate whether the MI analysis captures nonlinear relationships between the envelope and the neural response on the individual channel level. And third, we apply both measures to a group study (i.e., different age groups) for which we know significant group differences exist (Gillis et al., 2022), with the purpose of showing subtle yet important differences between both measures.

#### Temporal and spatial patterns

The linear Temporal Response Function (TRF) models the brain’s response to the stimulus, hence polarity naturally occurs. The TMIF, on the other hand, is a non-negative metric of neural tracking at each timestep. Nonetheless, the temporal and spatial patterns of both measures correspond well (see Figure 1). A negative peak in the TRF is also represented as a positive peak in the TMIF. Hence, TRF topographies with high positive (red) and high negative (blue) values are both high positive (red) in the TMIF topographies. The prediction accuracy in the linear model, which can be obtained by correlating the actual and the predicted EEG per channel, reflects neural tracking in the linear approach. The topography of these correlations resembles the MI (see Figure 1C).

We quantified the degree of correspondence between the two measures for the temporal patterns and neural tracking. For the temporal patterns, we computed the correlation between each corresponding channel in the TMIF and the absolute value of the TRF at the single-subject level. We used an integration window of 0 to 500 ms, including the peaks and latencies of interest. The group results are visualized in Figure 4A. Each datapoint represents the mean Spearman correlation of all channels for a single participant. Across participants, correlations are moderate to strong, with an average of r=0.57.

**Figure 4.**
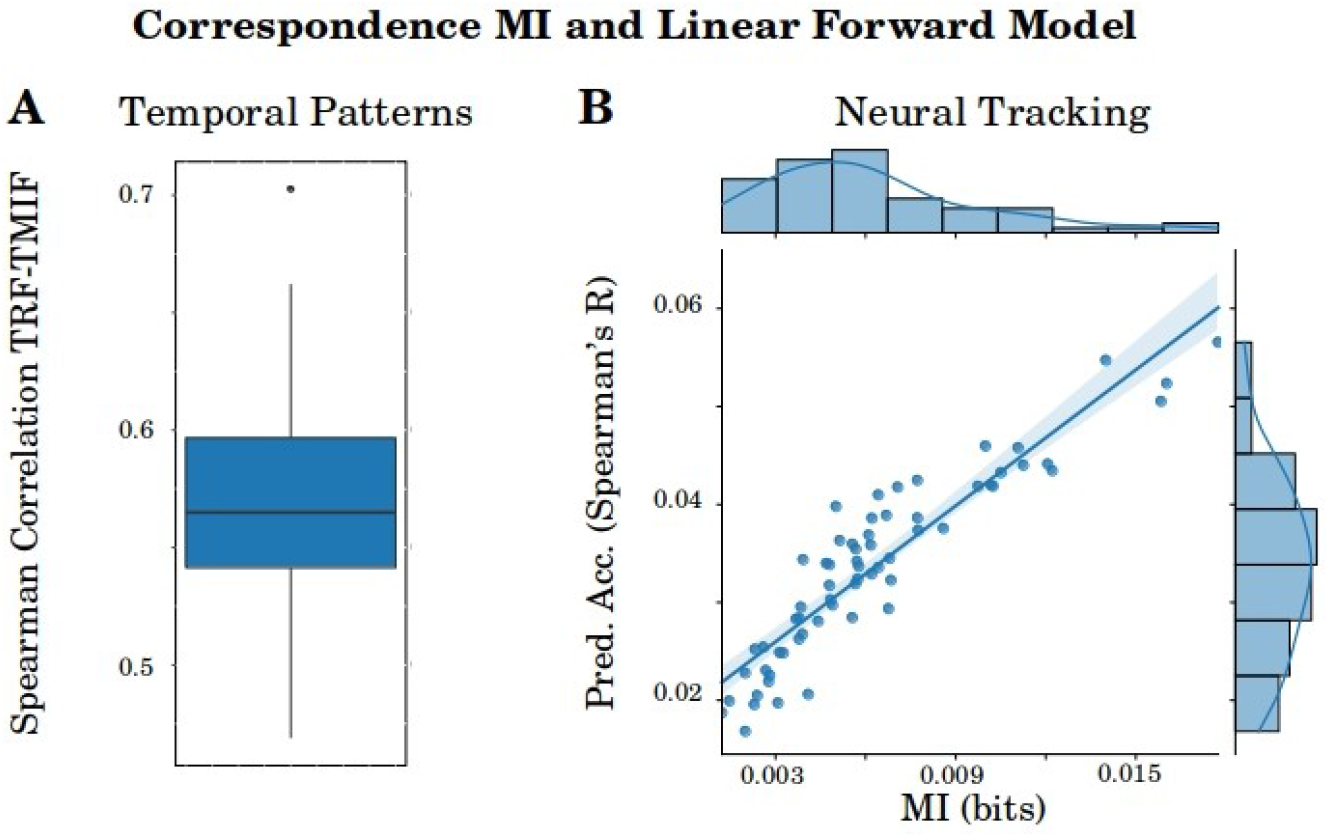
Correspondence between MI and the linear forward model. **A.** Correlations between all corresponding channels of the TMIF and the absolute value of the TRF in a 0 to 500 ms integration window. Each datapoint represents the mean correlation across all channels of one participant. **B.** Scatter plot of the mean MI and neural tracking in a 0 to 500 ms integration window. Each datapoint represents a single MI and Spearman’s R value. A regression was fit at the group level and the shaded area represents the 95% confidence interval.

The prediction accuracy quantifies neural tracking in the linear forward model, resulting in a single value per participant. To compare this measure to MI, we extracted the mean MI over the same integration window as the linear model, i.e., 0 to 500 ms. Both neural tracking measures were averaged over frontal and temporal channels. We fitted a regression over all participants (see Figure 4B) and report a Spearman correlation of r=0.94 (p<0.001). In conclusion, the MI and the linear forward model exhibit great correspondence.

#### Beyond linear components?

To check whether MI captures nonlinear components in the single-channel response to the speech envelope, we first filtered out the linear components in the data by subtracting the predicted from the actual EEG (see Methods). The TMIF between the residual EEG and the speech envelope was then calculated and interpreted against significance.

The TMIF for our channel selection is depicted in Figure 5A. Although the temporal pattern exhibits a reasonable evolution over time (i.e., with peak MI around 120 ms post-onset and a left-lateralized parietal topography), the TMIF did not reach significance on the group level. Not a single participant exhibited significant tracking of nonlinear components for the mean in a 0 to 400 ms integration window (Figure 5B). For the peak MI value, the TMIF reached significance in 27 out of 64 participants. The magnitude of MI for solely nonlinear components dropped with a factor of ≈ 1000 compared to the analyses including both linear and nonlinear components. This shows that, although there is significant envelope tracking for some participants for nonlinear components in the data, the added value of MI over the linear forward model is small and, in this case, not significant on the group level.

**Figure 5.**
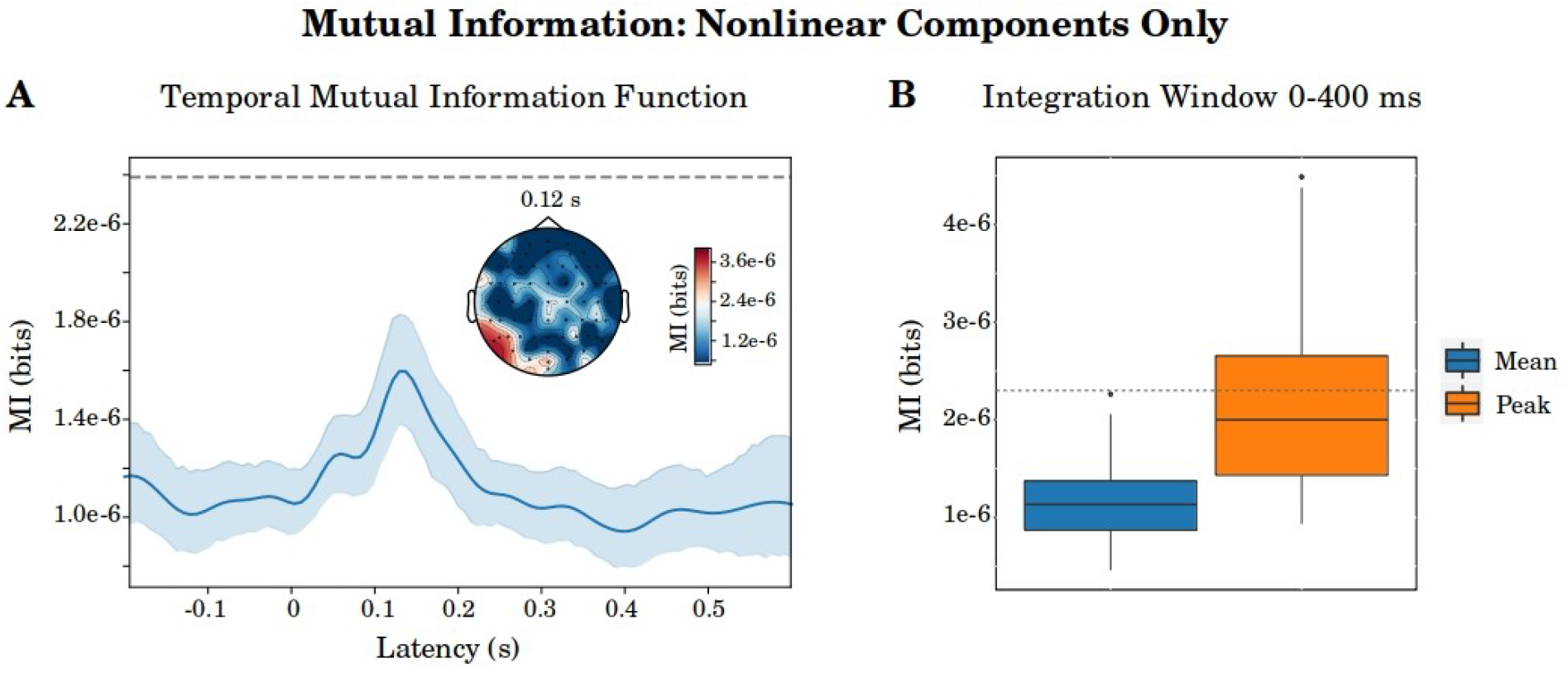
MI of nonlinear components. **A.** The TMIF between the speech envelope and the residual EEG (where linear components were removed from the EEG), and topography for the peak MI at 120 ms. The shaded area represents the 95% confidence inteval. **B.** Mean and peak MI in a 0 to 400 ms integration window. For both panels, the dashed grey line indicates the significance level.

#### Application to group study: healthy aging

Finally, we compare the linear forward model to MI in a group study. In a recent comparison, Gillis et al. (2022) found higher neural envelope tracking for younger listeners compared to older adults. We replicated these results (Wilcoxon signed-rank test, W=95, p=0.039, Figure 6A). Yet, enhanced neural tracking for younger listeners was not represented in the linear TRF amplitudes (a non-parametric cluster-based permutation test found no significant higher amplitudes for younger adults, Figure 6A). This is in contrast to the TMIF, where group differences do emerge in the spatio-temporal domain (a left parieto-occipital, p=0.007, and a right temporo-central, p=0.019) and on average over channels and latencies (W=107, p=0.005, Figure 6B).

**Figure 6.**
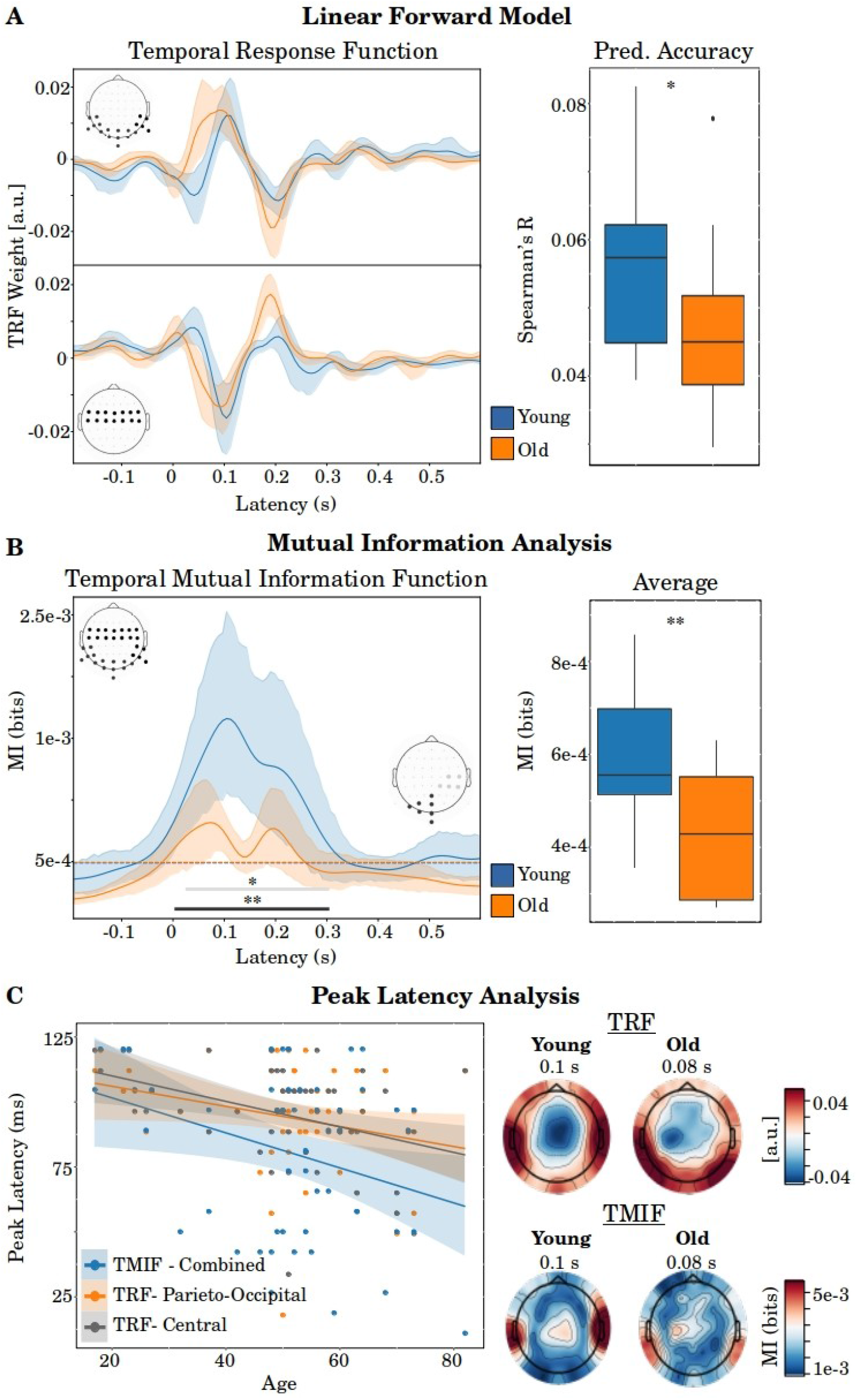
Application to group study. **A.** Results for the linear forward model. The TRF for the fronto-central and parieto-occipital channels are depicted, and the boxplots on the right represent the prediction accuracies for both channel selections combined in a 0 to 500 ms integration window. **B.** Results for the MI analysis. The TMIF for the combined channels is visualized, with bold horizontal lines indicating significant group differences (the same color is used to show the significant channels). Boxplots on the right represent the average in our channel selection in a 0 to 500 ms integration window. **C.** Peak latency analysis. The latency of the first peak shifts as age increases, captured by both methods. The topographies on the right show the associated peak topographies in the TRF and the TMIF. For all panels, the shaded area indicates the 95% confidence interval

A popular feature of the linear TRF is that it allows for peak latency analyses. In this dataset, a shift in the latency of the first peak was observed as age increased (Gillis et al., 2022). We plotted this trend for the parieto-occipital (positive peak) and central (negative peak) channels in Figure 6C (Pearson’s r=-0.27, p=0.048; r=-0.39, p=0.004, respectively). The correlation in peak latency is also present for the TMIF (channels combined; Pearson’s r=-0.31, p=0.020). In conclusion, we show that the TMIF can handle peak latency analyses with statistical relationships comparable to the TRF.

### 3.3 Multivariate Mutual Information in Relation to the Linear Backward Model

In the linear backward model, the speech envelope is reconstructed from a weighted sum of multiple EEG channels shifted in time. Traditional MI measures are restricted in their application to multidimensional data because they suffer from the curse of dimensionality (i.e., the number of parameters increases exponentially with the dimensions). The Gaussian copula MI has a particular advantage in that perspective since the number of parameters increases only linearly with the number of dimensions. Here, we apply this method to determine the multivariate relationship between the envelope and the EEG for Dataset 1. Figure 7A depicts the TMIF between the envelope and the 64-dimensional EEG data (64 channels combined). The MI peaks around 100 ms and reaches significance at expected latencies (+-0 to 300 ms). We further report strong correspondence between the mean multivariate MI and the linear backward model, both in a 0-500 ms integration window (Spearman’s r=0.93, p<0.001, Figure 7B).

**Figure 7.**
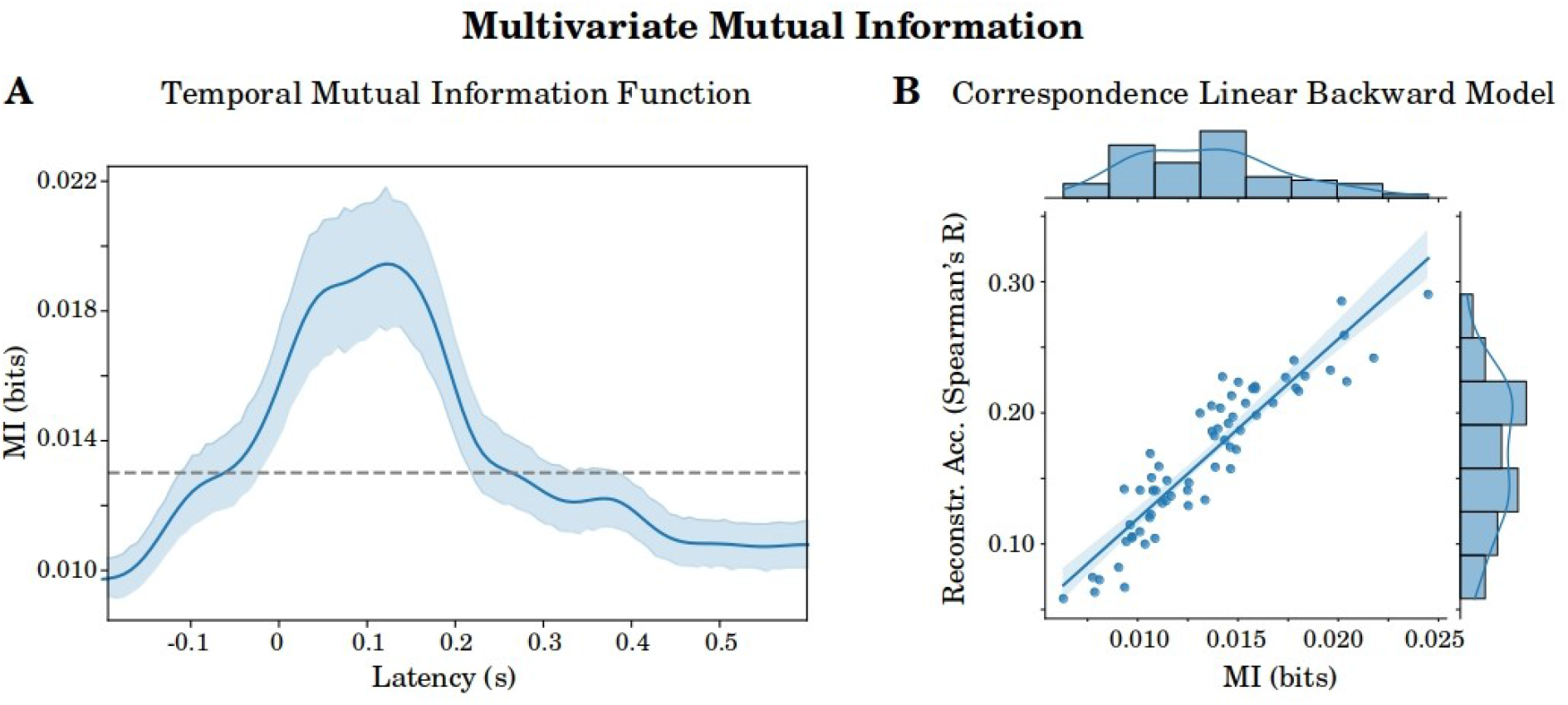
Multivariate MI. **A.** The TMIF for the multivariate MI, including all 64 channels. The grey line indicates the significance level. **B.** Correspondence between the multivariate MI and the linear backward model. Both methods used a 0 to 500 ms integration window (mean in the integration window for MI). Each datapoint represents a participant, the shaded area indicates the 95% confidence interval

#### Beyond linear components?

Section 3.2 reported that the single-channel MI analysis did not capture significant nonlinear relationships with the envelope. Here, we investigate whether such relationships do arise for the multivariate MI. Linear components captured by the backward model were removed by subtracting the reconstructed envelope from the actual envelope. Next, we calculated the TMIF between the residual envelope and the 64-dimensional EEG data.

The results are depicted in Figure 8. The temporal pattern reaches significance at the group level and peaks at 120 ms post envelope (Figure 8A). The mean MI (integration window 0 to 400 ms) is significant for 56 out of 64 participants, and the peak value reaches significance for 62 participants. Compared to the analysis that included linear components, MI dropped by a factor 10. This drop is largely reduced compared to the MI for single channels, where a factor of 1000 was observed (Section 3.2). We conclude that multivariate MI has significant added value over the linear backward model. Furthermore, we conclude that multivariate MI analysis is a more powerful tool for detecting nonlinearities in the data than single-channel MI analyses.

**Figure 8.**
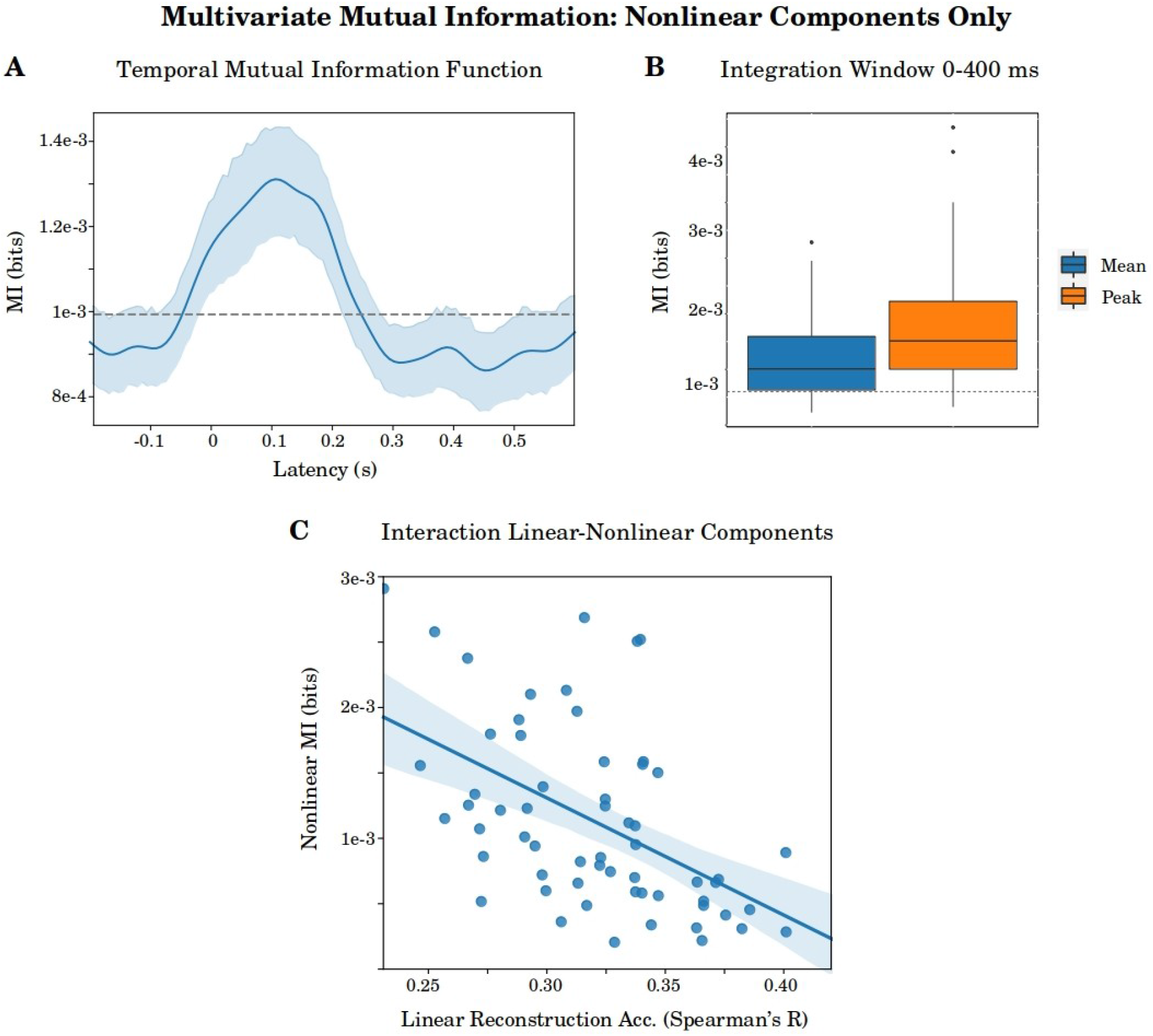
Multivariate MI of nonlinear components. **A.** The TMIF between the EEG and the residual speech envelope (where linear components were removed from the envelope). **B.** Mean and peak MI in a 0 to 400 ms integration window. **C.** Interaction between linear and nonlinear components. Scatter plot between the reconstruction accuracies obtained by applying the linear backward model and the nonlinear MI (MI between the EEG and the residual envelope, after removing linear components). Each datapoint represents a participant. Dashed grey lines indicate the significance level, shaded areas indicate the 95% confidence interval.

An interaction with linear relations can partially explain the spread across participants in the nonlinear MI. Figure 8C illustrates a negative correlation between linear and nonlinear relations in the data. Lower reconstruction accuracies for the linear backward model are strongly related to higher nonlinear MI values (Spearman’s r=-0.56, p<0.001). This shows that nonlinear relations between the EEG and the envelope tend to be more strongly represented when linear relations are less prominent and demonstrates the added value of a model that captures all kinds of variance in the data: there is no loss of information with nonlinear methods.

## 4 Discussion

The present paper provides an overview of the different methodologies to calculate MI for the study of neural envelope tracking. A comparison between the most commonly applied histogram-based approach and the Gaussian copula MI (Ince et al., 2017) showed that the former method is biased by the power spectrum of the EEG and the envelope. Furthermore, multivariate applications (e.g., neural tracking between the envelope and multiple channels combined) are limited since the histogram-based approach suffers severely from the curse of dimensionality. In contrast, the Gaussian copula MI does not suffer from these issues. The temporal evolution and spatial pattern of the Gaussian copula MI resembled linear models. The MI analysis also allows for peak latency analyses, with statistical relationships comparable to the linear forward model. This was demonstrated in a group study comparing younger and older listeners. Beyond linear components captured by the linear model, the MI analysis detected significant nonlinear components in the data. The effects were most strongly present for the multivariate MI. Finally, an interaction between linear and nonlinear relations was reported: participants with low neural tracking of linear components had relatively high tracking of solely nonlinear components. This shows that there is more nonlinear variance in the EEG when there is relatively less linear variance present. This nonlinear variance can only be captured when applying nonlinear techniques. Therefore, we argue that the MI analysis is a more informative tool for studying neural envelope tracking than linear models.

### 4.1 Comparing different Mutual Information Derivations

Section 3.1 showed that the histogram-based MI is biased by power in the low frequencies of the EEG and the envelope. The bias is present for (1) a baseline period where the brain has not yet processed speech and (2) permutation tests, where either the EEG or the envelope is reshuffled while the original power spectrum is retained. This shows that the bias is independent of the true relationship between the EEG and the stimulus. Further, we showed that with increasing bins, the histogram-based MI increasingly suffered from the bias (see Figures 2 and 3). The number of estimated parameters increases quadratically with the number of bins (400 vs 16 parameters for 20 vs 4 bins). As a result, errors in the joint probability estimations are more likely to occur, and a bias can become more prominent. We therefore strongly advise keeping the number of bins low.

The prominence of the bias is strongly data-dependent. A bias can become more prominent with few data and a relatively great difference in power in the low frequency oscillations across channels or stimuli. Yet, the question remains what bin parameter is optimal for a specific case. One could argue to specify the number of bins based on the difference between MI and the significance level. In this study, the maximal difference was found at 4 bins on the group level (see Figure 2E). However, with this approach, the bin parameter is selected post-hoc. This can lead to a research bias, and it can pose a challenge for group comparisons (when the optimal number of bins differs between groups). Maximizing MI relative to the significance level at the single-subject level is also not an option. With a higher number of bins, MI increases, which would make results non-comparable across subjects (seeFigure 2 and Vinh et al. (2009)). We thus lack an objective criterion to specify the optimal number of bins.

The Gaussian copula MI directly solves this issue as it does not require parameter values. Furthermore, the Gaussian copula MI is robust to a bias induced by low frequency power. In this approach, the number of estimated parameters (i.e., the covariance matrices) stays constant with the number of variables (2, in this case). Therefore, errors in the joint probability distribution are less likely to occur. The Gaussian copula MI has several other advantages over the histogram-based approach. First, the Gaussian copula approach can be used to calculate the multivariate MI between the envelope and a combination of channels. This contrasts the histogram-based approach where it is practically impossible to calculate the multivariate MI since the number of estimated parameters grows exponentially with the number of dimensions or variables. Second, as discussed in the introduction, erroneous high MI values cannot occur with the Gaussian copula method whereas the histogram-based approach can overestimate the true MI. Finally, previous work has shown that the Gaussian copula MI has higher sensitivity (i.e., a higher rate of correctly detecting a true relationship) compared to the histogram-based approach (Ince et al., 2017). For these reasons, we argue that the Gaussian copula method is the preferred MI approach to study neural envelope tracking.

#### Methodological considerations for mutual information applications

The MI between the EEG and the envelope is a relative measure and requires permutation testing. Although the bits scale provides a meaningful effect size (Friston, 2012), many factors can impact the magnitude of MI. These include the length of the stimulus and the EEG recording (shorter length results in higher MI), the sampling rate (lower sampling results in higher MI), and the power spectrum (higher power in lowest frequency bands result in higher MI by chance). Permutation testing must be performed by computing a null distribution that controls for these covariates. Random permutations of EEG- or envelope-shaped noise (shaped with the same power spectrum as the original signal) are both valid choices (Crosse et al., 2021; Nichols and Holmes, 2002). The 95th percentile of permutations can be taken as the alpha = 0.05 significance level. With the TMIF, individual time points can then be compared to the significance level. In line with our expectations, brain latencies following the envelope were significantly tracked while latencies preceding the envelope were not (see Figures Figure 1, 6 and Figure 7). Note that permutation tests are also required for linear methods (Crosse et al., 2021; Combrisson and Jerbi, 2015).

As discussed, the Gaussian copula MI can be applied to either individual EEG channels or a combination of channels (i.e., the multivariate MI). The former approach has the advantage that it allows for an interpretation of the spatial distribution of MI. In contrast, the latter method is more informative as it determines the multivariate relationship between the envelope and all EEG signals combined. Note, however, that there will be an upper limit to the number of dimensions (i.e., channels) that one can use in the multivariate MI, which will strongly depend on the amount of data. Future work needs to address the interplay between the amount of data and the maximum number of dimensions. In the present study, the envelope was significantly tracked for a combination of 64 channels with 15 minutes of data. We suggest that researchers relate the obtained outcome to the significance level. When neural tracking is not significant with a certain number of dimensions, researchers can consider applying data reduction techniques such as Denoising Source Separation (DSS) (Särelä and Valpola, 2005) or a Multiway Canonical Correlation Analysis (MCCA) (de Cheveigné et al., 2019). A Deep Canonical Correlation Analysis (DCCA) (Andrew et al., 2013) may be the preferred option, as it preserves the nonlinear nature of the data.

### 4.2 Mutual Information in Relation to Linear Models

The spatial and temporal patterns of the linear forward model (TRF) and the MI (TMIF) showed great correspondence. The TRF models the EEG response and retains polarity, while the TMIF reflects a measure of statistical dependency between the EEG and the envelope and is strictly positive. Both negative and positive peaks in the TRF were positive in the TMIF. Latencies and topographies of the peaks showed good correspondence. Across participants, the TRF and the TMIF correlated r=0.57 on average (see Figure 4A). Neural tracking measures for the linear forward (prediction accuracy) and backward model (reconstruction accuracy) were strongly associated with the MI in the single-channel and multivariate case, respectively. We report a correlation of r = 0.93 for both cases (see Figures 4B and 7B).

We applied the linear forward model and the MI analysis to a group study comparing envelope tracking for young and older listeners (see Figure 6). With both methods, we reported higher envelope tracking for young listeners. Yet, the TRF amplitudes did not reflect higher envelope tracking. By contrast, the TMIF exhibited higher MI magnitudes at the peak latencies for younger listeners.

An important mention when interpreting the group study results is that the TRF must not be interpreted as a measure of neural tracking. The TRF reflects the amplitude weights at different response latencies to predict the EEG. It is an arbitrary unit sensitive to scale differences. Therefore, it requires normalized data for interpretations (Crosse et al., 2021). Yet, the TRF does not determine the statistical relationship between the EEG and the envelope. To obtain precise temporal information on envelope tracking, researchers can apply narrow integration windows or fit separate models per latency (for applications of these approaches, see Vanthornhout et al. (2019) and Gwilliams et al. (2020), respectively). By contrast, the TMIF displays the statistical relationship between the EEG and the envelope at each response latency. The TMIF is insensitive to scale differences. It thereby provides precise temporal information of neural tracking. A single measure of neural tracking can be obtained by taking the average or maximum in the TMIF. Alternatively, one can calculate the multivariate MI between the envelope and a combination of response latencies in an integration window (I(EEG latency 1, EEG latency 2,…, EEG latency N; envelope), with N latencies).

In the linear forward model, the entire TRF waveform is optimized to predict the EEG. By contrast, the TMIF concatenates individually calculated MI values at each latency. As a result, the TRF has sharper aligned peaks and suffers less from autocorrelation than the TMIF. This is an advantage of the TRF. Another difference between both methods is that linear models require regularization and cross-validation to prevent overfitting. On the other hand, the proposed MI algorithm provides a direct statistical test for the dependency between the EEG and the envelope and does not require those techniques. Hence, there is no loss of data (i.e., no validation and test set), which is a significant added value of the MI analysis.

Recent advances in linear forward and backward models have investigated the neural response to speech features beyond the acoustic envelope. One particular feature that improves the models’ performance and gained much interest over the years is discrete phoneme categories (Di Liberto et al., 2015). The MI between a continuous (EEG) and a discrete (phoneme category) variable can also be calculated under the Gaussian copula framework. Here, separate probability distributions are fit per discrete value. For a more in-depth explanation, we refer to Ince et al. (2017). In a recent study, Daube et al. (2019) successfully applied the Gaussian copula MI to neural tracking of phoneme categories.

Assessing the added value of a feature in an MI framework can be done similarly to linear model comparisons. Researchers can subtract MI values between the EEG and a combination of features with a held-out combination. The obtained difference is then mathematically equal to the conditional MI (Cover and Thomas, 1991), *I*(*X*; *Y|Z*) (where X is equal to the feature set of interest, Y the EEG data, and Z the held-out combination of features). The conditional MI is the information-theoretic analog of a partial correlation. Researchers can apply it to unrelated sets of variables as well, where one assesses the effect of a variable while controlling for a third. Conditional MI was recently applied in the field of envelope tracking (Bröhl et al., 2022). Note, however, that the Gaussian copula approach has an upper limit on the number of dimensions one can use for accurate estimations of the covariance matrices. Future research needs to address this question.

Linguistic processing of natural speech is commonly investigated using sparse, continuous stimulus features. As an example, word surprisal or word frequency, a pulse aligned with the onset of a word, can improve the prediction accuracy (Gillis et al., 2021). Such sparse stimulus features may pose a challenge for the Gaussian copula MI. Many zero-entries will complicate the rank-step in the MI algorithm. One possibility is to use the feature value for the entire duration of a word or convolve it with a kernel. However, this requires further investigation. When using sparse stimulus features or for applications in high dimensional spaces, linear models or deep neural nets are the preferred options.

For phase data, the MI analysis has an advantage over linear models. The nonlinear MI analysis can capture the circular nature of phase data. This enables phase entrainment analyses (Cogan and Poeppel, 2011; Gross et al., 2013), phase-amplitude coupling (Combrisson et al., 2020; Keitel et al., 2018) and network-based statistics (Giordano et al., 2017). This opens avenues to more comprehensive and holistic analyses of the neural response to the speech envelope.

In summary, the MI analysis is a valid technique to study neural envelope tracking. It exhibits great correspondence with linear models in terms of neural tracking and spatio-temporal characteristics of speech processing. The MI analysis has the advantage that statistical dependency and temporal information are captured in the same measure, i.e., the TMIF. By contrast, the TRF can not be interpreted as a measure of neural tracking. Furthermore, the MI analysis does not require regularization or cross-validation of the outcomes, which must be applied when using linear models. Given the nonlinear nature of MI, researchers can apply the measure to phase data as well. On the other hand, linear models have an advantage in a large number of dimensions or when using sparse stimulus features (beyond the acoustic envelope only).

### 4.3 Beyond Linear Components

We investigated whether the MI analysis detects nonlinear components in the neural response to the envelope. Linear components were removed by applying linear models. Next, the TMIF for the residual data was calculated. The single-channel TMIF did not significantly track nonlinear components on the group level. However, for a substantial number of participants (27/64), we found a significant peak in the nonlinear TMIF. Furthermore, a reasonable evoked response with peak topography on the group level was obtained (see Figure 5). Therefore, we argue that a meaningful trend is present for neural tracking of nonlinear components. Nonlinear effects may be stronger for a larger amount of data or MEG data (which has a better signal-to-noise ratio).

For the multivariate case, the MI analysis detected significant nonlinear components on the group level (see Figures 8A and 8B). The multivariate approach accounts for interactions between channels which are precluded in the single-channel MI analysis. For this reason, the multivariate MI is a more powerful tool for detecting more complex relationships between the neural signal and the envelope. We further showed that MI values for solely nonlinear components were negatively correlated with linear relationships captured by the backward model (see Figure 8C). A relatively poor linear fit can thus partially be explained by the fact that linear models fail to detect nonlinear relationships in the data. Therefore, we conclude that the MI analysis is a more informative approach to neural envelope tracking.

Our results are in line with studies applying nonlinear deep neural networks. Better performance for deep neural networks compared to linear models have been reported in the past (Accou et al., 2021; Ciccarelli et al., 2019; de Taillez et al., 2020; Monesi et al., 2020; Vandecappelle et al., 2021; Xu et al., 2022). Yet, previous studies applied these models to different protocols, namely auditory attention decoding or a match-mismatch classification paradigm, and were often trained subject-independently. For an identical protocol, Thornton et al. (2022) recently demonstrated that reconstruction accuracy (decoding the speech envelope from EEG data) for subject-dependent deep neural networks is higher compared to linear models. Here, we replicate these results: neural envelope tracking benefits from techniques that capture both linear and nonlinear relationships. Compared to the MI analysis, deep neural networks can model more complex relationships with many more input features. A downside of deep neural networks is that they are black boxes and preclude spatial and temporal interpretations of speech processing. Such interpretations are retained when applying the MI analysis.

## 5 Conclusion

The Gaussian copula-based MI analysis is a valid alternative to linear models for studying neural envelope tracking. Researchers can apply this method to single-channel data and a combination of channels for both phase and amplitude data. The single-channel MI analysis enables spatial and temporal analyses of the neural response to the speech envelope. Beyond the limits of linear models, the MI analysis detects significant nonlinear components in the data. Effects are strongest when jointly considering all EEG channels. We conclude that MI is a statistically more powerful tool for studying neural envelope tracking while it retains spatial and temporal characteristics of speech processing.

## Acknowledgments

The authors would like to thank Bernd Accou and Lien Decruy for data sharing, and Marc Van Hulle, Alexander Bertrand and Robin Ince for their advice on the analyses.

## 6 Supplementary Figures

### Channel selection

**Supplementary Figure 1.**
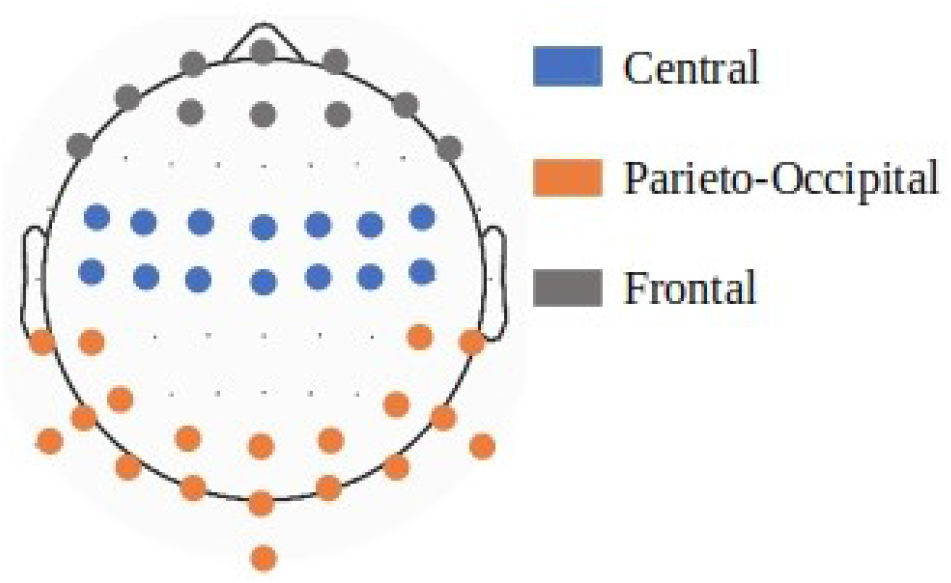
Display of the different channel selections, referred to in the Methods section.

### Example of biased results in the EEG

**Supplementary Figure 2.**
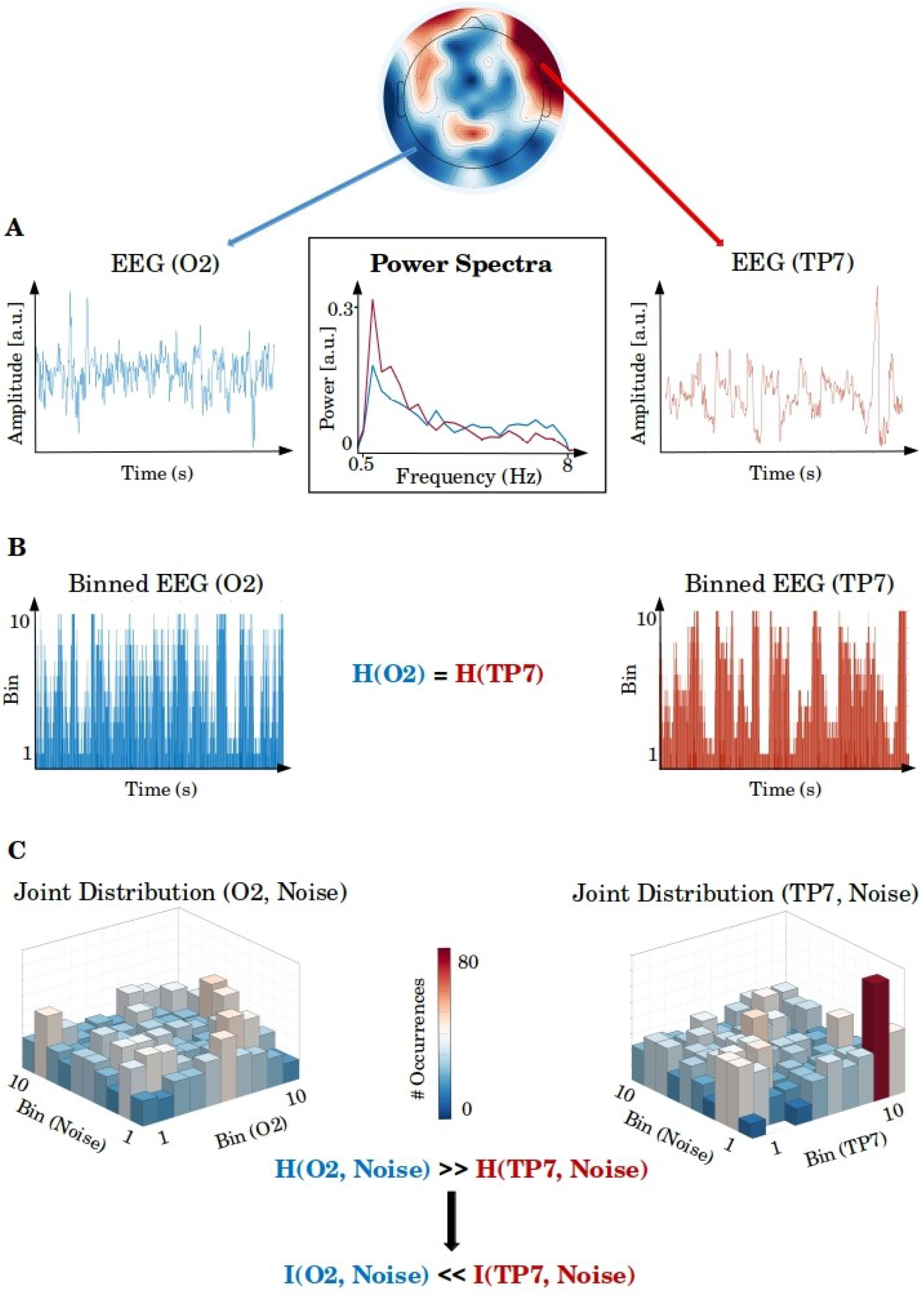
Example of bias present in frontal channels for 30 seconds of EEG data for 1 participant. **A.** EEG data for channel TP7 (frontal) and channel O2 (occipital) and their power spectra. **B.** Binned representation of the corresponding EEG data. Entropy for both channels are equal since the equipartition principle is applied. **C.** Joint distribution between the EEG and the speech envelope. The joint distribution for channel O2 is approximately uniformly distributed, while transient jumps in bin combinations occur for channel TP7. Consequently, channel TP7 has lower joint entropy and higher MI.

## 1 Appendix A

### 1.1 Calculating Mutual Information

Mutual Information (MI) between two variables X and Y is expressed as a function of the marginal entropies and the joint entropy:

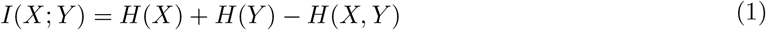

with

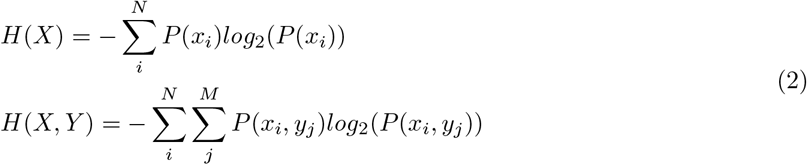

Where H(X) is equal to the entropy of the marginal distribution of variable X (e.g., the speech envelope), and *P*(*x_i_*) the probability of observing value *x_i_* with N possible values. The entropy of a second variable, Y (e.g., the neural response of a single channel), is computed according to the same formula. The joint entropy is represented by H(X,Y) and expresses the uncertainty in the joint distribution of X and Y.

Temporal information of MI (i.e., the TMIF) in the context of neural envelope tracking is obtained by sliding the neural response in a channel (Y) in function of the envelope (X) for a certain integration window. A schematic overview of the MI and the procedure to obtain the TMIF is represented in Figure 1.

**Figure 1.**
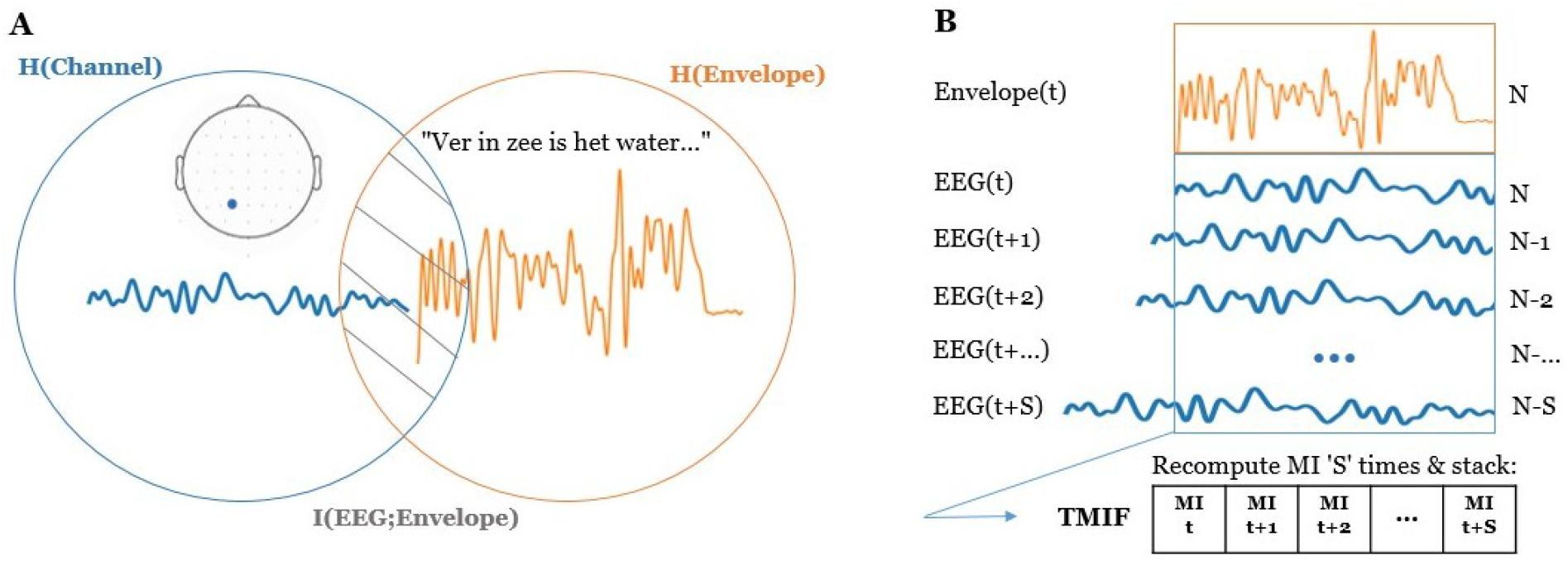
**A.** Conceptual representation of Mutual Information (MI) in the context of neural envelope tracking using a Venn diagram. MI is the amount of shared information between the EEG and the speech envelope, obtained through subtracting the sum of the individual entropies with the joint entropy. **B.** The Temporal Mutual Information Function (TMIF). The TMIF is obtained through shifting the EEG in function of the envelope ‘S’ times and recomputing MI at each timestamp.

In order to determine the MI between the EEG and the envelope, we require the observation probabilities of the individual datapoints. This must be estimated from the data. As outlined in the introduction, several alternatives exist to construct the probability distributions. Here, we explain the methods in more detail.

#### The histogram-based method

The most straightforward approach to estimating the probability distribution is based on the histogram-based algorithm. Here, the data are first partitioned into a set of discrete bins. The probability of observing a discrete value is obtained by dividing the number of occurrences in a single bin by the total number of data points. The probability values can then be plugged into the formulas of marginal and joint entropy.

The binned approach is appealing due to its simplicity. However, there are several considerations to take into account. First, the data discretization can be performed via two main strategies: either bins are set with equal bin width (‘equidistant’), or bins are set to have equal occupancy (‘equipartition’, e.g., if using 10 bins, each bin contains 10% of the data). The equidistant approach estimates the true probability distribution, while the equipartition binning transforms the data to a uniform distribution.

The equipartition approach is the preferred option for two main reasons. First, it is robust to sampling errors. In the equidistant approach, bins with low occupancy can receive too low probability due to insufficient sampling, not reflecting the true probability curve. In contrast, the equipartition binning assigns equal observation probabilities to all bins. And second, the equidistant approach facilitates MI comparisons over variables since the entropy for all individual variables is equal for a certain number of bins (i.e., all variables have uniform distributions). Hence, the MI will not depend on the marginal entropies, but will be fully encapsulated by the joint entropy (see Eq. 1). This is preferred as the joint entropy determines the relationship between both variables. By contrast, the distributions and marginal entropies differ across variables in the equidistant approach. As a result, the MI is a complex interplay between the marginal and the joint entropy.

A second consideration is the number of bins to choose for constructing the histogram. In theory, one should not have too few bins, otherwise, there will be little sensitivity to the distribution, and the relationship between two variables can be lost. On the other hand, if one has too many bins, estimation errors for the joint distribution can be made hence generality is lost. The latter case can be seen as a form of overfitting, as the distribution will become sensitive to noise components in the data. Therefore, the bin parameter can induce either an under-(too few bins) or overestimation (too many bins) to the true MI. Previous neural tracking studies have chosen the bin parameter rather arbitrarily with little justification or empirical data to support their choice.

#### Continuous alternatives

Instead of partitioning the data into discrete bins, several continuous alternatives exist to estimate the probability distribution. In the Kernel Density Estimation (KDE) approach, each datapoint is convolved with a (most commonly Gaussian) kernel. The probability density function is then obtained by summing all kernel values and normalizing so the integral equals one. In the KDE approach, researchers need to pre-specify the width of the kernel (i.e., the bandwidth), which is the equivalent critical choice to the number of bins in the histogram-based approach (Venelli, 2010). The bandwidth parameter determines the smoothness of the obtained probability distribution. A bandwidth set too large leads to over smoothing, hence specificity is lost, while a bandwidth too small leads to under smoothing, hence generality is lost. A recent neural envelope tracking paper calculated MI based on KDE estimations (Keshavarzi et al., 2021), but the bandwidth parameter was not specified.

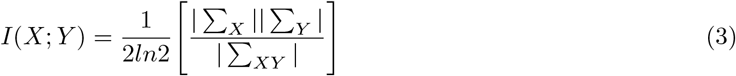

where ∑_*X*_ and ∑_*Y*_ are the covariance matrices of variables X and Y, and ∑_*XY*_ is the covariance matrix for the joint variable. Straight lines indicate that the determinant of the covariance matrices is calculated. The resulting algorithm does not require any parameter values. However, we must rely on the strong assumption that the data is normally distributed.

#### Gaussian copula method

Recently, Ince et al. (2017) introduced the concept of Gaussian Copula MI for neuroimaging research. The method uses the Gaussian definition of MI but transforms the data, so it meets the normality assumption. The procedure first implies that the individual variables are ranked on a scale from 0 to 1, obtaining the cumulative density functions (CDF’s). These ranked variables are then uniformly distributed. The joint distribution of these two variables, i.e., the joint CDF, is expressed in function of the marginal CDF’s and a function that links these two variables (Sklar, 1959). This function is called the copula, and it describes the statistical relationship between the two variables. The entropy of the joint CDF is equal to the sum of the entropy of the marginal CDF’s plus the entropy of the copula (Ma and Sun, 2011):

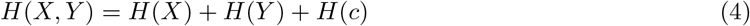

where H(c) is the entropy of the copula. When plugging this formula for the joint entropy into the general formula of MI (Eq. (2)), we obtain:

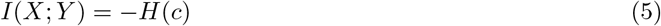

MI is equal to the negative entropy of the copula (Ma and Sun, 2011). We thus no longer require the entropy of the marginals, as the MI is fully captured by the copula (i.e., the statistical relationship between both variables). As a result, the MI can be calculated by estimating the entropy of the copula. Yet, this again results in the problem of accurately estimating probability values for the copula. Instead, Ince et al. (2017) suggest exploiting the fact that the MI does not depend on the individual variables (see Eq. (5)). Therefore, we can transform the marginal distributions in any way we see fit as long as the copula is preserved. By computing the inverse standard normal CDF of the variables separately, the data distributions are transformed to perfect standard Gaussians, and the copula is preserved. As a result, the parametric MI estimate in Eq. (3) can be applied.

In summary, the Gaussian Copula TMIF in the context of neural envelope tracking is obtained by applying the following steps:

1. Rank the speech envelope and the neural time series of a single channel on a 0 to 1 scale, obtaining the marginal CDFs as uniform distributions
2. Transform the marginal CDFs to standard Gaussians by computing the inverse standard normal CDF
3. Create the joint variable by stacking the envelope and the neural time series as a matrix sized time by 2
4. Compute the covariance matrices and calculate MI according to the formula in Eq. (3)
5. Repeat for all channels and time lags

The transformations of the data distributions are depicted in Figure 2.

**Figure 2.**
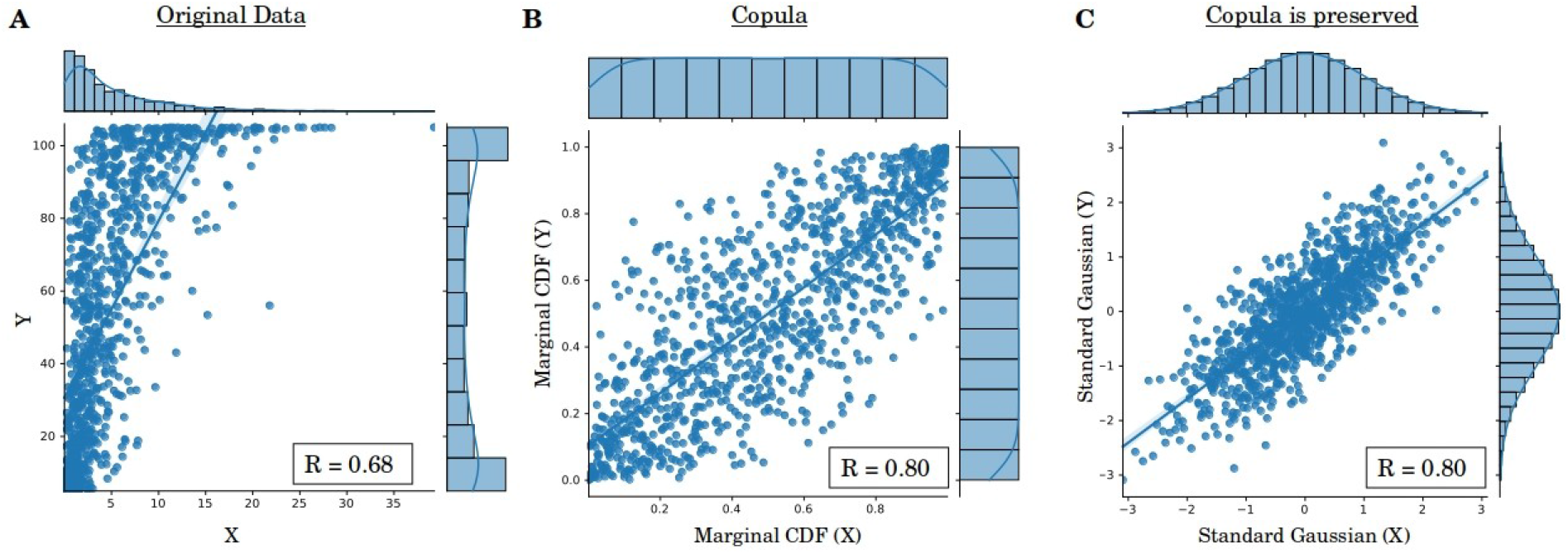
Data transformations applied in the Gaussian copula MI. **A.** Scatter plot of simulated data drawn from the Gamma (X) and Beta (Y) distributions. **B.** Data are first ranked on a scale from 0 to 1, resulting in uniform distributions. The joint distribution represents the copula. **C.** By taking the inverse standard normal CDF, perfect standard Gaussian marginals are obtained but the empirical copula linking both variables is preserved.

This algorithm is computationally efficient and makes no assumptions on the distributions of the variables. Crucial to mention, the MI in this method cannot be overestimated, i.e., erroneous high values cannot occur. This is because the joint Gaussian distribution has the maximum entropy for a variable with a given mean and covariance matrix (Cover, 1999). As MI is the negative copula entropy, the MI derived from a Gaussian copula is a lower bound to the true MI (Calsaverini and Vicente, 2009). For a more in-depth explanation of Gaussian copula MI, we refer to Ince et al. (2017). Over the past recent years, the Gaussian copula MI has been applied in numerous neural envelope tracking papers (Coopmans et al., 2022; Daube et al., 2019; Giordano et al., 2017; Perez et al., 2022).

#### Multidimensional Variables

The Gaussian copula has a particular advantage over the other MI derivations in multidimensional spaces. While the histogram- and KDE-based approaches severely suffer from the curse of dimensionality and computational complexity, the Gaussian copula method can efficiently handle multidimensional variables. After transforming each individual dimension to a standard Gaussian, the formula in Eq. (3) can be applied. For the context of neural envelope tracking, this may open perspectives to calculating the MI between the speech envelope and multiple channels combined: I(Envelope; Channel 1, Channel 2,… Channel N). This could be considered an information-theoretic analog to the linear backward model.

